# VirTect: a computational method for detecting virus species from RNA-Seq and its application in head and neck squamous cell carcinoma

**DOI:** 10.1101/272278

**Authors:** Atlas Khan, Qian Liu, Xuelian Chen, Yunjing Zeng, Andres Stucky, Parish P. Sedghizadeh, Daniel Adelpour, Xi Zhang, Kai Wang, Jiang F. Zhong

**Affiliations:** Department of Medicine, Columbia University, New York, NY, 10032, USA; Raymond G. Perelman Center for Cellular and Molecular Therapeutics, The Children’s Hospital of Philadelphia, 3501 Civic Center Boulevard, 5th Floor CTRB, Philadelphia, PA 19104; School of Dentistry, University of Southern California, Los Angeles, CA 90033, USA

**Keywords:** Viral detection, RNA-seq, Human papillomavirus (HPV) carcinogenesis

## Abstract

Next generation sequencing (NGS) provides an opportunity to detect viral species from RNA-seq data on human tissues, but existing computational approaches do not perform optimally on clinical samples. We developed a bioinformatics method called VirTect for detecting viruses in neoplastic human tissues using RNA-seq data. Here, we used VirTect to analyze RNA-seq data from 363 HNSCC (head and neck squamous cell carcinoma) patients and identified 22 HPV-induced HNSCCs. These predictions were validated by manual review of pathology reports on histopathologic specimens. Compared to two existing prediction methods, VirusFinder and VirusSeq, VirTect demonstrated superior performance with many fewer false positives and false negatives. The majority of HPV carcinogenesis studies thus far have been performed on cervical cancer and generalized to HNSCC. Our results suggest that HPV-induced HNSCC involves unique mechanisms of carcinogenesis, so understanding these molecular mechanisms will have a significant impact on therapeutic approaches and outcomes. In summary, VirTect can be an effective solution for the detection of viruses with NGS data, and can facilitate the clinicopathologic characterization of various types of cancers with broad applications for oncology.

**Significance Statement:** We developed a new bioinformatics tool, and reported the new inside of HPV carcinogenesis mechanism in HPV-induced head and neck squamous cell carcinoma (HNSCC). This novel bioinformatics tool and the new knowledge of HPV-induced HNSCC will facilitate the development of target therapies for treating HNSCC.

## Introduction

The advent of next generation sequencing (NGS) provides an opportunity to accurately and comprehensively detect and identify viral pathogens in clinical samples (1, 2). Researchers have developed some computational tools for the discovery and identification of viruses in NGS data from human tissues (3–5), including viral integration sites (6–10). However, all previous studies used subtractive analyses. For example, in a study conducted by Isakov et. al. (3), three steps were utilized for virus identification: (I) align short reads against a human reference genome using tophat (11), (II) subtract non-human sequences from human sequences, and then (III) categorize viruses based on nucleic acid databases. Also Kostic et. al. used a subtractive approach to detect viruses (4); they considered the reads which were not mapped to the human genome after filtrations to be “candidate nonhuman pathogen-derived reads.” Another set of researchers used a subtractive approach with thresholds for coverage and minimum number of reads, which were mapped to virus genomes in their defined virus database (5). They used two different filtrations: (I) the number of mapped reads to any virus (>1000) and (II) the minimum coverage of reads (30X). Using the aforementioned methods, it is possible to encounter some noise or poly(A) sequences which may also have high coverage and a high number of reads aligned to viral sequences; however, they do not actually represent a viral sequence. Thus, to address this issue, there is an urgent need to develop more robust computational tools which can correctly detect and identify viruses from human tissues, and discriminate viral sequences from background noise or sequencing artifacts.

Here, we describe a new method called VirTect for detecting viruses in RNA-seq data from human clinical samples. Our approach to virus detection is different from existing methods in that we include unique filters to discriminate real viral sequences from noise and artifacts, thereby minimizing false positive rates. We use three different types of filtration in VirTect: (I) a threshold for the number of mapped reads, (II) a threshold for the coverage of mapped reads, and (III) a threshold of the length of continuous mapped regions for any pathogen genome in our defined virus database. A non-human sequence which passes these three phases of filtration would very likely be a viral sequence, and by employing this method, VirTect can significantly improve the accuracy of detecting pathogenic viruses to inform clinical diagnosis, therapeutics, and prognosis.

In the present work, we demonstrate the utility of this bioinformatic tool by perfoming detailed analysis of cancer RNA-seq data from patients. With the awareness of chemical carcinogens such as tobacco, certain types of cancer have steadily decreased. However, cancers induced by viral infection have increased significantly (12–14). Recent studies indicate that approximately 12% of human cancers are viral in etiology (15). For example, human papillomavirus (HPV) comprises a group of more than 200 related viruses, and some HPV subtypes were found to be associated with specific types of cancers such as cervical, vaginal, vulvar, anal, penile, and head and neck squamous cell carcinomas (HNSCC) (16–18). The oncogenic HPV subtypes (*e.g.*, subtypes 16, 18, 31, 33, 35, 39, 45, 51, 52, 56, 58, 59 and 68) are sexually transmitted viruses (19–22). Importantly, the proportion of HNSCCs that are HPV-positive increased from 18% in 1973 to 32% in 2005 in the USA, representing an unprecedented and dramatic epidemiologic spike (23). However, the mechanism of carcinogenesis in HPV-induced HNSCC is not well studied compared to that of HPV-induced cervical cancers. In the present work, we performed detailed analysis of RNA-seq data from 363 HNSCC patients using VirTect, and identified 20 HPV16-positive and 2 HPV33-positive patients. The viral and host (patient) gene expression profiles suggest that the E2 and E7 genes of HPV, but not E6, play a major role in HPV-induced HNSCC. The molecular mechanism of HPV oncogenesis in HNSCC differs from what has been reported in cervical carcinogenesis(24). The viral/host gene expression profiles of HPV-induced HNSCC elucidate HPV oncogeneis in HNSCC. In doing so, this new bioinformatic tool provides actionable targets for developing new diagnostic assays and therapeutics for HNSCC. VirTect is available at https://github.com/WGLab/VirTect

## Results

An overview of VirTect is shown in Figure 1. NGS data in FASTQ format provided the input, which was then mapped to a human reference genome using TopHat (11). After the subtraction of human sequences, nonhuman sequences were aligned using BWA-MEM against the nonhuman sequences in our defined virus database (currently 757 different viruses) to identify viruses. Finally, VirTect performed filtrations to discriminate true viral sequences from noise or artifacts and reported identified virus(es).

**Figure 1:**
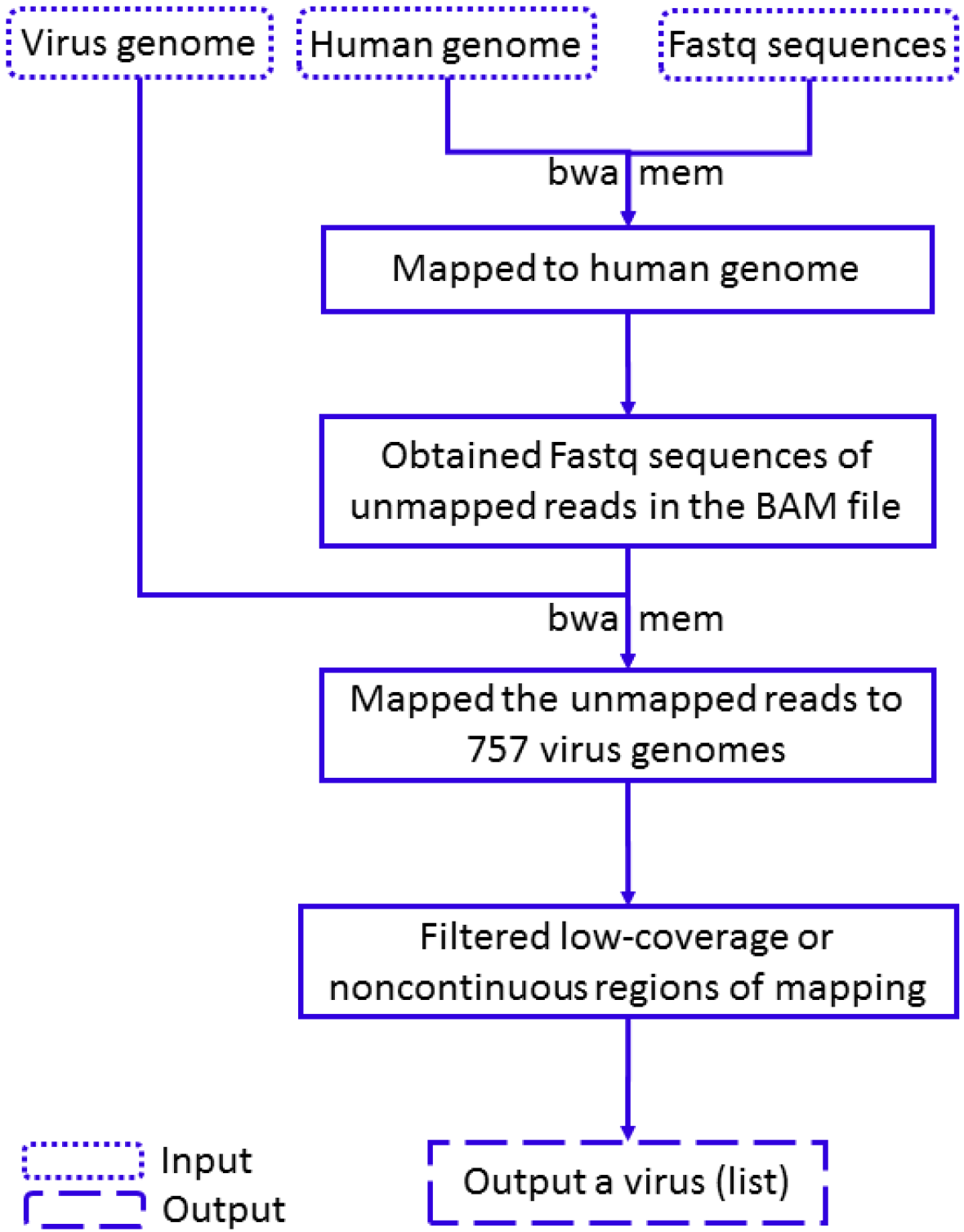
Schematic overview of the VirTect pipeline for virus detection from next generation sequencing (NGS) data.

### Detection of HPV in HNSCC patients with VirTect

To test the accuracy of VirTect, we analyzed 363 HNSCC samples acquired from the TCGA database and detected HPV transcripts in 22 cases, of which 20 contained HPV16 transcripts and 2 had HPV33 transcripts. We examined pathology reports as a validation of our predictions. Histopathology and clinical assays agreed with our RNA-seq analysis results (Table 1). The histopathology of H&E slides from each HPV+ case was rigorously examined by a pathologist to confirm the morphology of viral infection. Histopathology findings (detailed in Figure 2) confirmed morphologic features consistent with HPV infection and confirmed the HNSCC phenotype in each case. A random subset of 30 negative cases were also examined, and the results suggested that samples without detectable HPV genes were indeed HPV- pathologically.

**Figure 2:**
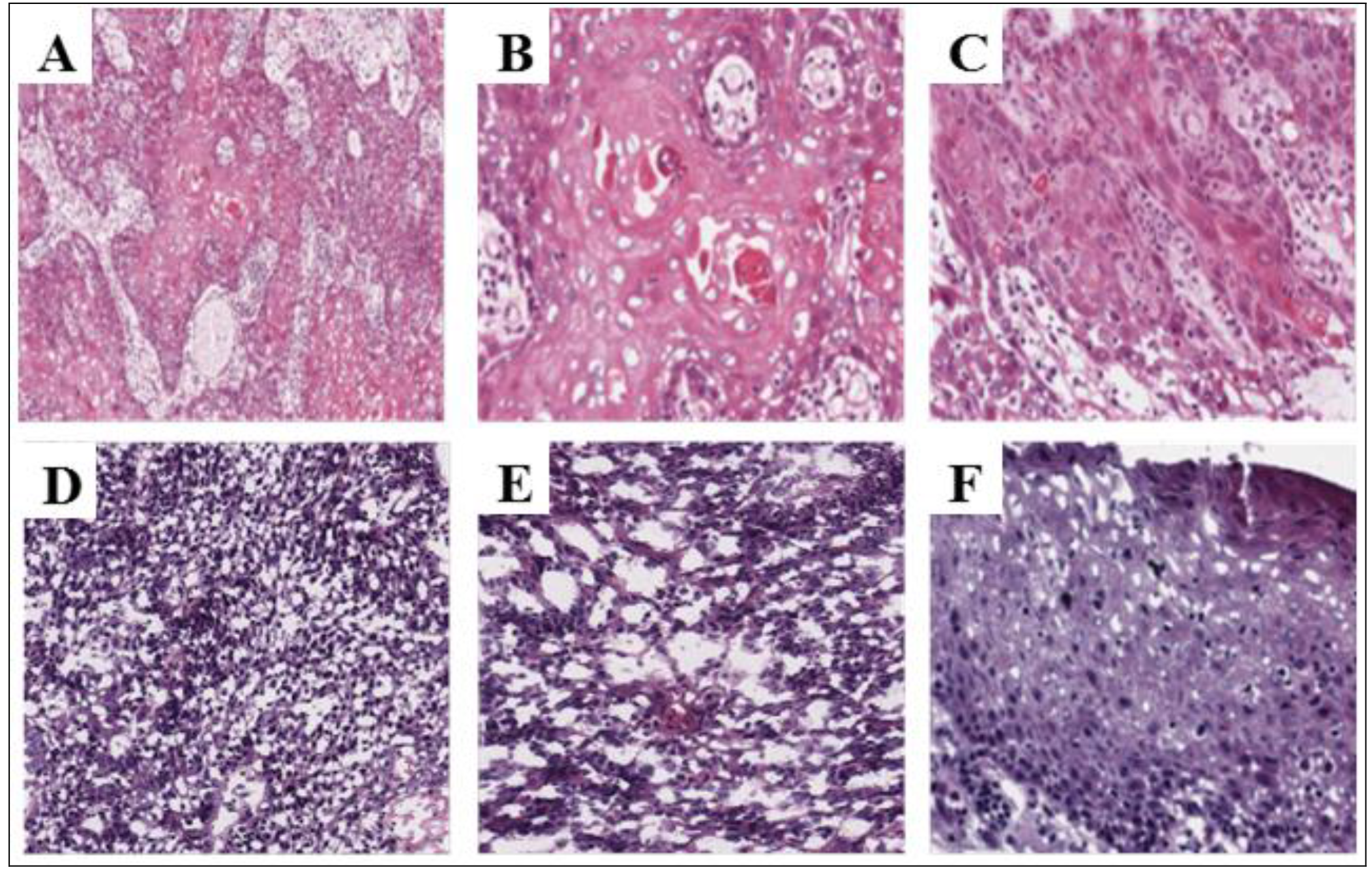
Histopathology confirms *in silico* results of HPV infection. **A-C.** Histopathology of cancer specimens without *in silico* HPV detection. **A.** Moderately differentiated invasive squamous cell carcinoma (20X). **B.** Higher magnification showing invasive tumor islands with central keratin pearl formation and individual cell dyskeratosis (40X) **C.** Cellular atypia, pleomorphism, nucleoli, and mitotic activity are observed regularly (40 X). **D-F.** Histopathology of *in silico* HPV-positive specimens which have characteristic non-keratinizing tumor morphology. **D.** A typical *in silico* HPV16-positive specimen (20X). **E.** Higher magnification (40x) shows infiltrating tumor islands lacking squamous maturation and comprising a cell population of ovoid to spindle-shaped cells with indistinct borders or gap junctions, and hyperchromatic nuclei that lack prominent nucleoli. **F.** Overlying mucosa and significant koilocytosis indicating HPV infection. Viral cytopathic effect can be seen in the form of keratinocyte nuclear enlargement and hyperchromasia with perinuclear clearing or halo.

**Table 1:** Clinical confirmation of HPV16 infection in HNSCC cases.

| Patient number<br>(TCGA Case Number) | Sample location | Pathological<br>Evidence | P16 Assay<br>Confirmation | Morphology<br>Confirmation | Sex |
| --- | --- | --- | --- | --- | --- |
| **1** ( abbcc1d8-ab74-4459-bf28-cc627bef440e ) | Oropharynx/Tonsil | Oropharynx/Tonsil | P16 positive<br>(cytoplasm and<br>nuclei) | Yes | Female |
| **2** (e37f479f-ca05-45e4-ac6d-a974bed6e7f8) | Left tonsil | Left tonsil | NA | Yes | Male |
| **3** (5ac57aee-4be1-4a29-a53f-343f5a3d2e86 ) | Pharynx,<br>oropharynx | Pharynx, oropharynx | NA | Yes | Male |
| **4** (cac30b32-03ef-4ecb-88d8-d5577ab6b2a6) | Right base of tongue | Right base of tongue | NA | Yes | Male |
| **5** ( 59a7d695-cb0c-47b8-ba9d-ca52ca2cfa7d) | Right tonsil | Right tonsil | P16 positive | Yes | Male |
| **6** (cd032ddb-55f2-4c77-8dcf-e4e630f7de6f) | Right tonsil | Right tonsil | NA | Yes | Male |
| **7** (05f01280-bf77-4682-a7a8-20dd0eac77bd) | Left tonsil and soft<br>palate | Left tonsil and soft<br>palate | NA | Yes | Male |
| **8** (c7df3466-b9a7-4818-883b-d0cd08483570) | Right tonsil | Right tonsil | NA | Yes | Female |
| **9** ( 4bfbce2b-9d0b-4e8a-950f-fd8e0ba3e05a) | Right base of the<br>tongue | Right base of tongue | NA | Yes | Female |
| **10** ( 387db1df-ebaa-41c2-b036-f46ad61e313a) | Base of the tongue | Base of the tongue | NA | Yes | Male |
| **11** ( 9d469689-0413-4898-aa83-c6756cbfe117) | Right soft palate | Right soft palate | P 16/18 in-situ<br>hybridization<br>positive | Yes | Male |
| **12** ( 54b295d4-6315-4416-bde7-a221859f965c) | Left glossotonsillar<br>sulcus extending<br>into lingual tonsil | Left glossotonsillar<br>sulcus extending into<br>lingual tonsil | NA | Yes | Male |
| **13** ( b3631718-9e0a-454c-bee1-8f36ebc509d8) | Left tonsil |  | NA | Yes | Male |
| **14** ( 27c28c89-f4e7-4aec-a806-c0da7756e47f) | Right floor of mouth | Right floor of mouth | NA | Yes | Male |
| **15** ( 03c3ae62-d0aa-412e-bd3c-4577fc9f919c) | Right tonsil | Right tonsil | Positive for p16<br>and HPV16. | Yes | Male |
| **16** ( 9f89510c-ed07-471f-b35e-7c87c237b9fe) | Left tonsil | Left tonsil | NA | Yes | Male |
| **17** ( 602f2512-00b6-44b6-9ed6-f8b010224f8c) | Pharynx,<br>oropharynx | Pharynx, oropharynx | P16 IHC<br>positive, P16<br>ISH positive | Yes | Male |
| **18** ( 180036a2-3b56-405e-a1fe-d5932517b6c7) | Right tonsil | Right tonsil | NA | Yes | Male |
| **19** ( 10f53522-9ae7-47b9-80aa-b4b481561465) | Right tonsil | Right tonsil | NA | Yes | Male |
| **20** ( a0b136fb-3a0a-4411-8907-4ca775c7d04e) | Left tonsil | Left tonsil | P16 staining<br>positive | Yes | Male |

To compare VirTect with existing tools, we ran VirTect, VirusSeq and VirusFinder on a RNA-seq data set, in which pathology reports are available to indicate whether a patient has HPV16 infection. We used the metrics of precision (i.e., Predicted HPV+ cases confirmed with pathology / Total predicted HPV+ cases), recall (i.e. Predicted HPV+cases confirmed with pathology / All pathological HPV+ cases) and accuracy (i.e., Predicted cases confirmed with pathology/Total cases) of virus detection for comparison (Table 2). The results demonstrated that all the three tools had a precision of 1.00. However, VirusSeq had much lower recall (0.087); VirusFinder had a recall of 0.913. That is to say, VirusSeq and VirusFinder could not identify all samples where HPV16 virus was found in the pathology reports. In contrast, VirTect demonstrated 100% consistency with the pathology reports. We recognize that this is a relatively small sample size due to limited availability of pathology reports; nevertheless, this analysis confirmed that VirTect compares favorably to competing approaches on clinical samples.

**Table 2:** Comparison of virus detection by VirTect, VirusSeq and VirusFinder.

| Method | Precision | Recall | Accuracy |
| --- | --- | --- | --- |
| VirTect | 1.00 | 1.00 | 1.00 |
| VirusSeq | 1.00 | 0.087 | 0.344 |
| VirusFinder | 1.00 | 0.913 | 0.938 |

Note: RNA-seq data was available for 32 cases, including 23 HPV16+ and 9 HPV16-. **Precision:** Predicted HPV+ cases confirmed with pathology / Total predicted HPV+ cases. **Recall:** Predicted HPV+cases confirmed with pathology / All pathological HPV+ cases. **Accuracy:** Predicted cases confirmed with pathology / Total cases.

HPV16 transcripts encoding key viral oncoproteins (i.e., E7, E4, E5_a and E1∧E4) were detected in most of the HPV-positive samples. The HPV gene expression in an HPV16+ case, as visualized in IGV, is shown in Figure 3 (a). In this sample, the HPV16 oncoprotein E7 was expressed with high coverage and continuously mapped regions. A HNSCC sample with detectable HPV33 viral genes is shown in Figure 3 (b). The key viral oncoprotein E7 was also expressed in this sample. The HPV+ samples we identified were all oropharyngeal in origin, which further supports the link between HPV infection and oncogenesis since nearly all oral HPV+ cases are oropharyngeal in origin anatomically. Pathology reports in some of these cases included in situ hybridization for HPV high-risk subtypes which indicated HPV 16 infection, providing further molecular support for viral etiology.

**Figure 3:**
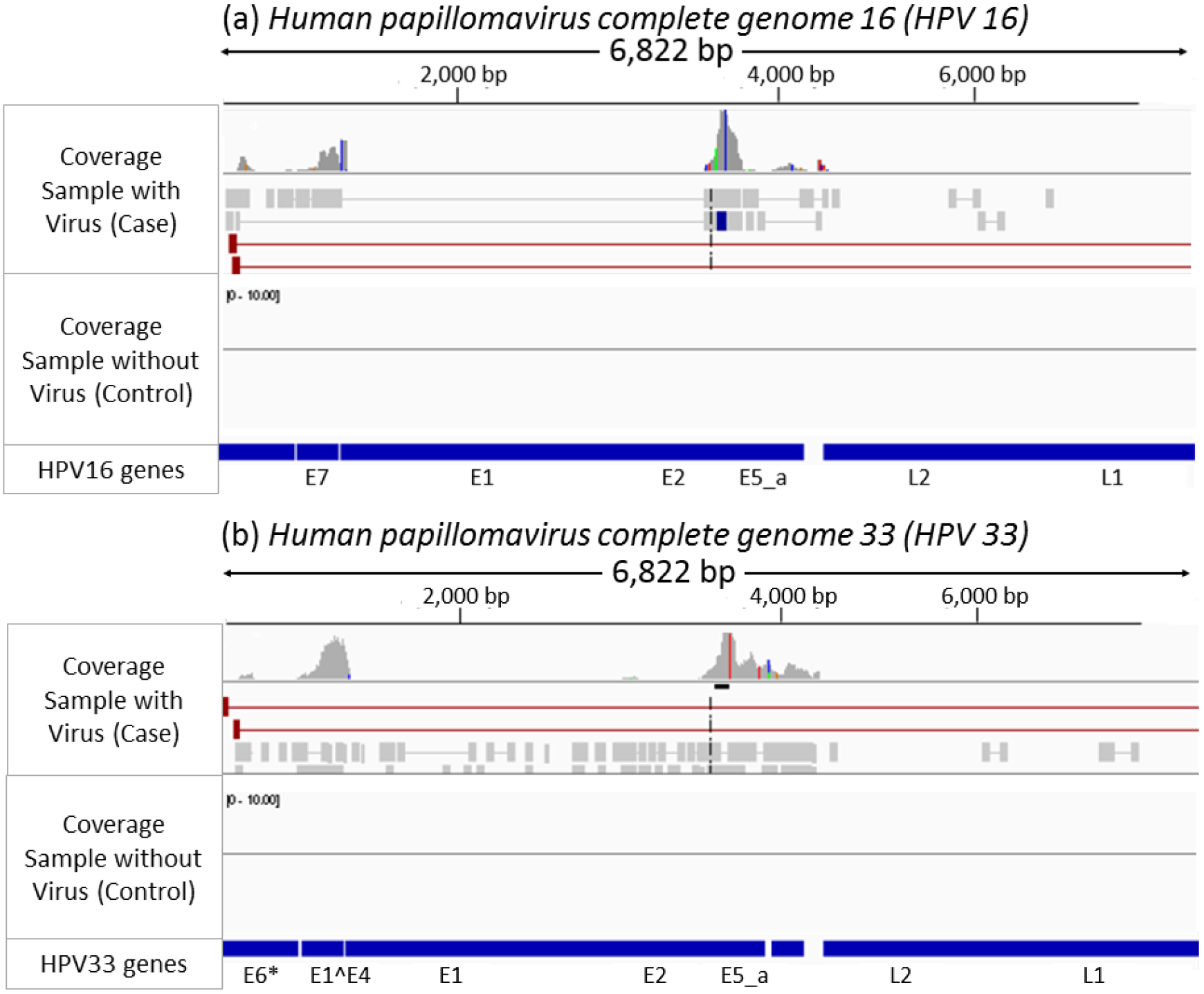
The IGV figures of the non-human reads of two head and neck cancer samples affected by HPV16 (a) and HPV33 (b) showing the coverage of a sample affected by HPV16/HPV33 and control.

Although pathology analysis is the gold standard for HPV infection, it is too labor-intensive for screening a large number of patients. With VirTect, we were able to screen 363 patients for HPV infection within a week. Among the 363 HNSCC patients, HPV 16 viral genes were detected in 20 patients (3 female and 17 male; see Table 1). Although the sample size is small, it appears that in this database, HPV-induced HNSCC was not equally distributed between males and females. Also, as expected, the L1 and L2 viral genes were not detected in any patient, and E2 and E8∧E2 were detected at high levels in all HPV16+ patients.

## HPV gene expression profiles in HNSCC

In cervical carcinoma, integration of the HPV16 viral genome into the host genome often leads to the disruption of the E1 and E2 open reading frame (ORF), resulting in unregulated expression of E6 and E7(25, 26). However, this does not appear to be the case in HNSCC (Figure 4). Among the 20 patients with detectable HPV16 viral sequences, the number of detectable viral genes ranged from 3 genes (patients 9, 11 &15) to 10 genes (patient 19). E2 and E7 were the most common, detected in 85% (17/20) and 90% (18/20) of patients, respectively. Most patients (15/20) had both detectable E2 and E7. The alternatively spliced E8∧E2 was detected in 90% (18/20) of patients, and the two patients (patients 9 and 15) without E8∧E2 were among those without detectable E2; these patients only had 3 detectable viral genes (E1, E1∧E4 and E7). E1∧E4 and E4 were detected in 95% (19/20) and 85% (17/20) of patients, respectively, in agreement with a previous report where E1∧E4 was found to be the most abundantly expressed HPV protein in infected epithelial cells(27). E6 (sequence range 83-559) was only detected in 30% (6/20) of patients. Fragments of this gene, namely E6*(sequence range 83-226) and E6*# (sequence range 409-417), were detected in 25% (5/20) and 15% (3/20) of patients, respectively. Patients 16 and 17 had detectable E6 and E6*, while all E6 fragments were detected in patient 19. This RNA-seq analysis suggested that both E2 and E7 were expressed at high levels in HNSCC. On the other hand, E6 was only detected in 30% (6/20) of patients, and the expression level was not as high as that of E7 or E2. In addition to HPV16 and HPV18, which have been reported to be causative agents of carcinoma(28, 29), we also found two HPV33-infected patients. The viral genes E7, E4, E1∧E4 and E8∧E2 were expressed in both patients, while HPV16 genes were not detected. These findings together suggest that HPV-induced HNSCC has a different molecular foundation from that of HPV-induced cervical carcinoma.

**Figure 4:**
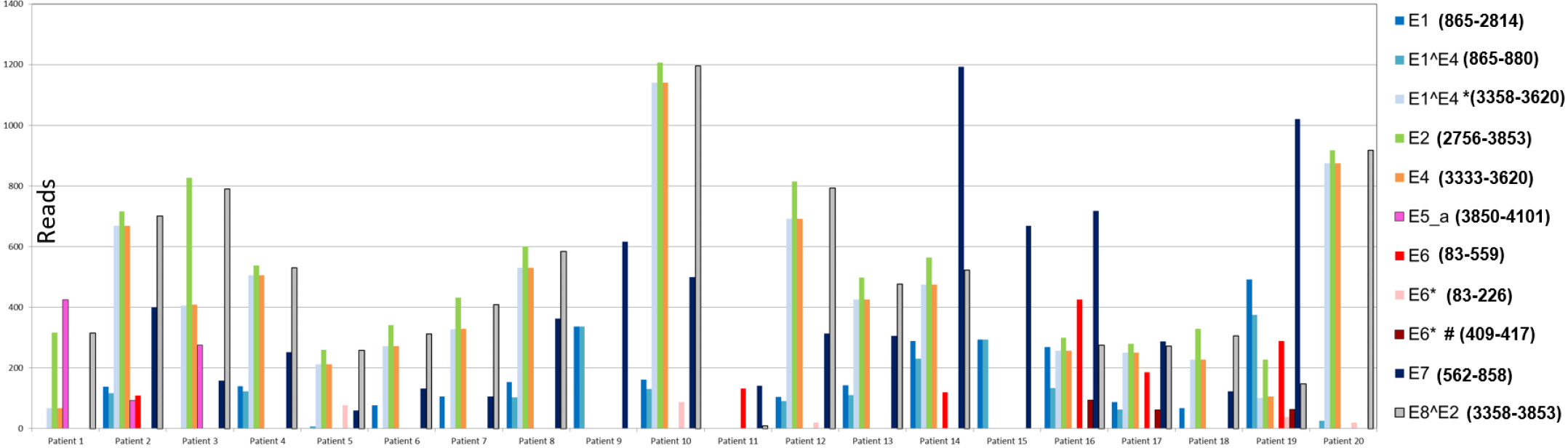
HPV16 viral genes detected in HNSCC patients. Among the 20 patients with detectable HPV16 viral sequence, the number of detectable HPV16 genes ranged from 3 (patients 9, 11 &15) to 10 (patient 19). E2 and E7 were most common, detected in 85% (17/20) and 90% (18/20) of patients, respectively. Most patients (15/20) had both detectable E2 and E7. The alternatively spliced E8∧E2 was detected in 90% (18/20) of patients. The two patients (patients 9 and 15) without E8∧E2 only had 3 detectable viral genes (E1, E1∧E4 and E7). E1∧E4 and E4 were detected in 95% (19/20) and 85% (17/20) of patients, respectively. However, E6 (83-559) was only detected in 30% (6/20) of patients. Fragments of E6*(83-226) and E6*# (409-417) were detected in 25% (5/20) and 15% (3/20) of patients, respectively. Patients 16 and 17 had detectable E6 (83-559) and E6*(83-226), while patient 19 had all E6 fragments detected. This viral expression pattern suggests that E2 and E7 are the major players in HPV-induced HNSCC.

### Molecular pathways in HPV16-induced HNSCC

Genetic instability is caused by steady accumulation of DNA damage and genetic variants, which lead to the activation of proto-oncogenes or the inactivation of tumor suppressor genes. A frequent event in cancers of the head and neck is the deletion of the short arm of chromosome 9, which results in inactivation of the host *p16* gene. In HPV-induced carcinogenesis, the most important initiating factor is the expression of the viral proteins E6 and E7, as they lead to the inactivation of the cellular tumor suppressor proteins p53 and Rb (30). The p16 expression assay is currently used clinically to identify HPV infection in HNSCC patients, because HPV+ individuals have very different prognosis and treatment options from those with non-virus-associated HNSCC. We identified highly significant upregulation of the tumor suppressor proteins CDKN2A (p16), CDKN2D (p19) and Tp53 in the samples from head and neck cancers with detected HPV16 infection when compared to samples without HPV16.

With Ingenuity Pathway Analysis, we identified a significant enrichment of proteins involved in the G1/S checkpoint in HPV+ samples (Figure 5). We first performed gene differential expression (DE) analysis by comparing 20 HPV16+ and 339 HPV16-HNSCC samples. We then used HTSeq for counting reads (31) and DESeq for detecting DE genes (32). We detected 437 genes with adjusted p<0.05 that showed significantly different expression levels between HPV16+ and HPV− groups. For DE genes, we performed enrichment analysis using DAVID (https://david.ncifcrf.gov/), which reported 22 pathways involving the DE genes that were differentially expressed between HPV+ and HPV− samples. The most probable pathway is illustrated in Figure 5, which shows how the G1/S cell checkpoint was altered in HPV16+ HNSCC patients when compared to HPV− samples. Several molecules in this pathway were differentially expressed, with large differences found in CDKN2D (p16), CDKN2A (p19), E2F1, and TP53 expression. Tp53 is the degradation target of E6, so a higher mRNA level of Tp53 suggests a compensation mechanism of P53 expression to offset the E6 degradation. The expression level of TP53 in the HPV+ HNSCC samples was slightly higher than that in the HPV-HNSCC samples, which correlated well with the low level of E6 detected. This result further suggests that the carcinogenesis of HNSCC is different from the mechanism established for the better-studied cervical cancer.

**Figure 5.**
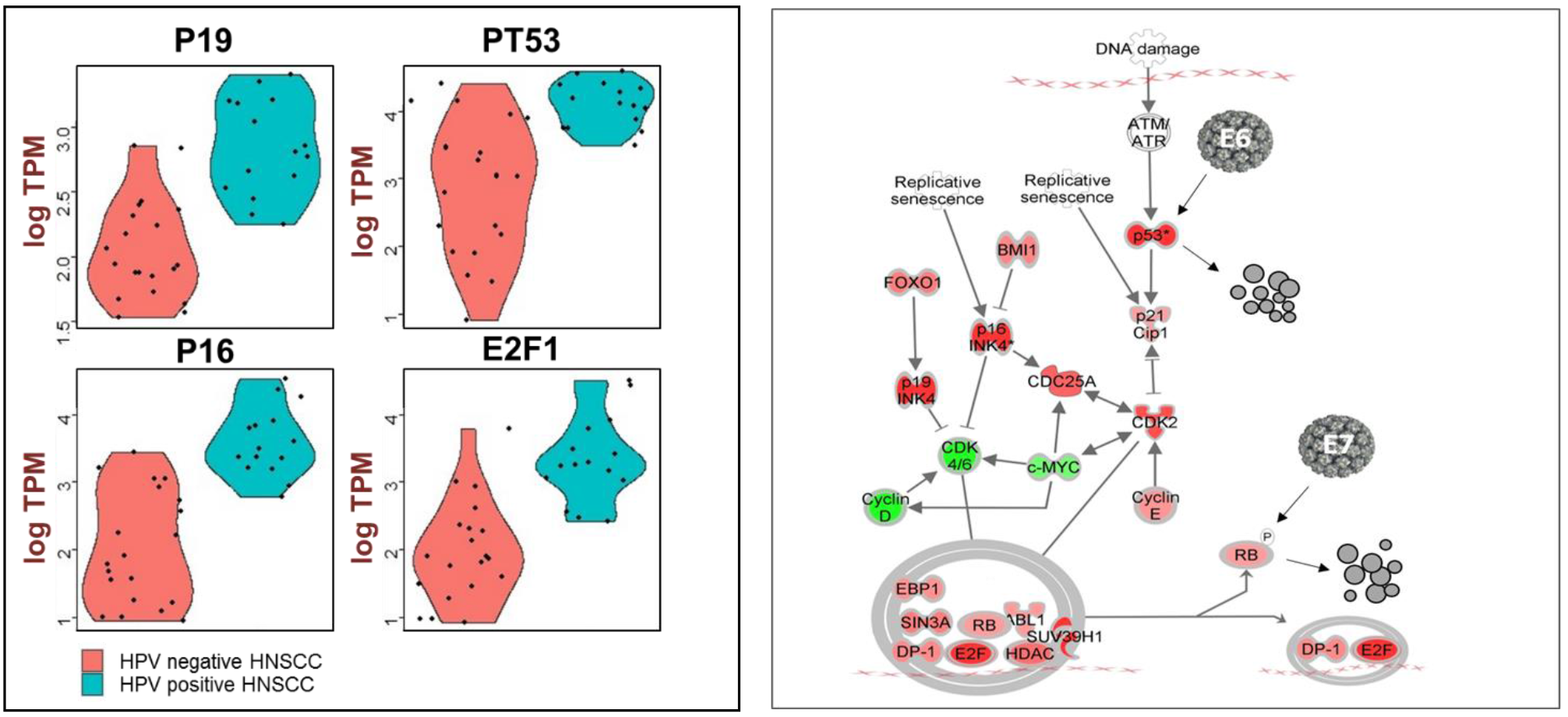
Carcinogenesis pathway of HPV-induced HNSCC. **A.**) HNSCC patients with (n=20) and without (n=20) HPV infection had significantly different expression values for p16 (p value=4.88E-7), p19 (p =7.6E-6), E2F1 (p =3.63E-5), and TP53 (p =9.75E-7). **B.**) Ingenuity Pathway Analysis (IPA) identified the molecules involved in the G1/S checkpoint to be the most significantly (p =3.28E-18) active molecular pathway, including 17 genes with significantly different (p <0.05) expression values between HPV+ and HPV− HNSCC. Red: upregulated in HPV+ HNSCC; green: down-regulated in HPV+ HNSCC. While E6 plays a minor role, E7 plays a major role in HPV carcinogenesis in HNSCC.

## Discussion

Here we reported a novel virus detection method called VirTect that uses RNA-seq data as input. We tested and analyzed VirTect’s performance on several datasets and identified several viruses from this analysis. Several groups have developed and discussed virus detection methods based on subtraction of nonhuman sequences from human samples (3–5). For example, VirusSeq (5) uses RNA-seq as input, and performs two filtrations to distinguish viral and non-viral sequences: the minimum number of reads aligned to a viral sequence (set at 1000 reads) and the coverage of the aligned reads (threshold of 30X). However, in some cases, a sample has thousands of nonhuman sequences, which may have repeat sequences that align to a virus but do not represent real viral genomes. Also in some cases, this approach may fail to detect real viral sequences, since coverage can be less than 30X; this may occur when viral infection is in latency stage and the coverage is low. Therefore, we sought to develop a new method that would detect viruses from NGS data with greater sensitivity and accuracy, by designing a new multi-layered filtering strategy. To confirm that this approach performs better, we compared VirTect with existing virus detection methods, namely VirusSeq (5) and VirusFinder (7). We found that VirTect could more accurately detect and discriminate noise and artifacts from real viral sequences. For example, for one head and neck cancer sample, VirusSeq wrongly attributed a poly(A) sequence to Hepatitis C and tick-borne encephalitis, again due to the high coverage (more than 30X) and more than 1,000 reads mapped to these viruses; however, it did not detect the real HPV16 sequences in this sample, since the number of mapped reads to HPV16 was 926, just below the threshold. In comparison, VirTect detected both the poly(A) sequence and the real HPV16 sequences (Table 1 and Figure 3); after filtration, VirTect indicated the presence of HPV16.

With VirTect, we also analyzed 363 HNSCC samples available in the TCGA database. We detected HPV in 22 of these samples; 20 had detectable HPV16 genes and 2 had detectable HPV33 genes. We confirmed the viral etiology of these cancers by correlating RNA-Seq and VirTect data to clinical and histopathologic findings. HPV-mediated carcinogenesis is thought to work through viral oncogenic proteins E6 and E7 via the disruption of cell cycle regulatory components(33). Specifically, E6 directly binds P53 and complexes it with the E6-AP ubiquitin ligase. This ubiquitination targets P53 for proteolytic degradation(34) and impairs host cell G2/M checkpoint and apoptosis, resulting in uncontrolled proliferation and genomic instability(35). Similarly, HPV E7 targets retinoblastoma (Rb) for degradation(36), and results in a deregulated cell cycle and unstable genome. However, our data suggest that E7 plays a major role in the carcinogenesis of HNSCC, while E6 only plays a minor role. The E7 viral gene was detected in all 22 HPV+ HNSCC samples, but E6 was only detected in 6 of the 22 patients (20 HPV16+ and 2 HPV33+). Our findings are supported by a recent study of HPV16-driven oropharyngeal squamous cell carcinoma (OPSCC), which showed that HPV16 E6 seropositivity has low sensitivity (50%, 95% CI19-81) but is highly specific (100%, 95% CI 96–100) (37).

To our surprise, E2 was detected in almost all HPV+ HNSCC patients. For cervical cancer, it has been reported that the integration of the HPV genome into the host genome often interrupts the E2 promoter and leads to unregulated E6 and E7 expression (25, 26). E2 is not detected in most cervical cancer cell lines with HPV infection(38). Furthermore, re-introduction of E2 to cervical cancer cell lines and HeLa cells was found to represses E6/E7 expression and lead to cellular senescence and apoptosis (39–43). In our study, E2 and E1∧E4 were detected at high levels, which agreed with the previous report that E1∧E4 stabilizes E2 (44). The detection of E2 and E8∧E2 viral fragments in RNA-seq data from HNSCC patients suggested that HPV-induced HNSCC has different molecular mechanisms of carcinogenesis from those of HPV-induced cervical cancers. Also, the cervical cancer high-risk strain HPV18 was not detected in any of the tested samples, but HPV33 was detected. Although only two patients had detectable HPV33 fragments, this is consistent with the report that HPV33 rarely induces HNSCC(45).

In light of these findings, we compared the gene expression profiles of HPV+ and HPV− HNSCC to identify a potential molecular pathway of HPV carcinogenesis in HNSCC. Our results agreed with the previous report that not all HPV16 infections associated with HPV16 E6 seropositivity(46). This underscores that HPV16 E6 seropositivity is not a suitable assay for HPV-induced HNSCC screening. Importantly, we presented evidence here that, although cervical and oropharyngeal mucosal cancers are generally thought to have the same mechanisms of carcinogenesis given that they are associated with similar HPV subtype infections (e.g., HPV16), in fact these two anatomical sites may experience different paths to carcinogenesis, even with the same HPV16 subtype. This potentially has profound translational implications for diagnostics and therapeutics in head and neck oncology.

We tested and evaluated VirTect on different datasets from TCGA, and we believe that VirTect can accurately detect viruses in NGS samples with greater accuracy than other methods. The ability to identify viral infection accurately may have important translational value for oncology, as we have demonstrated with a HNSCC dataset. This work has significant clinical relevance because of the importance of viral identification and characterization for precision medicine. For example, head and neck cancers which are associated with HPV16 have a better prognosis and are more sensitive to radiation therapy than non-HPV cancers. Therefore, accurate detection and knowledge of HPV status is essential for diagnosis, targeted therapeutic approaches, and prognostication, and ultimately improves patient outcomes (47). The main limitation of VirTect is that it depends on a database of all known viruses, which is used to nominate candidate viruses in human cancer tissue. Though the database is extensible, this approach cannot detect novel viruses that are not in the database. In our future studies, we will extend VirTect to exome and genome sequencing data for virus detection. We believe that VirTect will also be highly useful for analyzing whole-genome sequencing data.

## Materials and Methods

### Datasets

We evaluated and tested VirTect on 363 samples of head and neck squamous cell carcinoma (HNSCC) which were downloaded from the Cancer Genome Atlas (TCGA) (https://cancergenome.nih.gov/).

### Quality control and trimming

We performed quality control (QC) using FASTQC https://www.bioinformatics.babraham.ac.uk/projects/fastqc/, and then used cutadapt http://cutadapt.readthedocs.io/en/stable/guide.html to trim paired end reads of all samples to ensure that the quality was appropriate for further investigation. We did the following filtrations using cutadapt version 1.2.1 [12]:

I. Removed adaptor contamination, which may affect virus detection.
II. Trimmed the reads when the average quality score in a sliding window fell below a phred score of 20.
III. Discarded reads shorter than 40 bp.

The cutadapt command line was as follows:

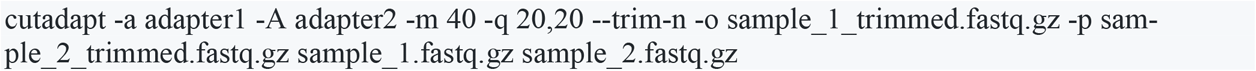

where *adapter1* is a forward adapter, *adapter2* is a reverse adapter, *-q* is for quality and *-m* is used for the minimum length of reads (i.e., reads less than 40 bp long were trimmed in our analysis). After trimming of all samples, we used VirTect for virus detection.

### Virus detection method (VirTect)

VirTect contains several steps, as illustrated in Figure 1:

**Step 1**: Paired end (PE) reads are aligned to a human genome reference to subtract the nonhuman sequences from the whole-genome sequence of human samples using TopHat (11).
**Step 2:** The nonhuman sequences derived in the first step are subtracted from the human sequences. In the second step, the bam file of the nonhuman reads is converted to FASTQ format using bedtools (48).
**Step 3**: The nonhuman sequences are mapped to the 757 different viruses currently in the database using bwa-mem (49) and the number of reads is quantified according to the mapping to the viruses in the defined database.
**Step 4:** Filtrations are performed to remove noise/artifacts or poly(A) sequences, which may have high coverage with thousands of reads mapped to virus genomes but may not represent real viral sequences. Three types of filtrations are used for this purpose:

I. Empirical cut-off of the number of the reads mapped to virus genome. A default cut-off is 500 in this study.
II. Cut-off of the coverage of reads. A default setting is 5X, i.e., coverage will be greater than 5.
III. Cut-off for continuously mapped regions. Our threshold is 100 by default but this is a user-predefined parameter. Since we used 48 bp sequences, this worked well for the data analyzed here; however, when reads are longer (more than 100bp or larger), this threshold will need to be increased.
**Step 5:** Finally, a list of viruses is generated from the samples that passed the filtrations, and this list can then be subjected to further investigation.

## Availability of data and materials

Our implementation of VirTect is available as a software tool at https://github.com/WGLab/VirTect. VirTect was implemented in Python programming language and has been tested on Linux platforms. It depends on third-party publicly available tools, including Samtools (50), bedtools, Bowtie 2, TopHat and BWA.

## Author Contributions

AK developed methods, analyzed data, interpreted data and wrote the manuscript. QL contributed to method development, data analysis and manuscript revision. KW and JZ coordinated the study. PPS and XZ provided clinical samples, evaluated and interpreted the data, prepared the pathology figures and data, and edited the manuscript. All authors discussed the biological findings and read/approved the final version of the present manuscript.

## Acknowledgments

The results published here are in part based upon data generated by The Cancer Genome Atlas (TCGA), managed by the NCI and NHGRI. We are grateful to TCGA for this source of data. Information about TCGA can be found at http://cancergenome.nih.gov. We also thank the Wang lab members for helpful comments and feedback on VirTect. This work was supported by NIH grants CA197903 (J.Z.) and HG006465 (K.W.).

